# Feasibility of spiral fMRI based on an LTI gradient model

**DOI:** 10.1101/805580

**Authors:** Nadine N. Graedel, Lars Kasper, Maria Engel, Jennifer Nussbaum, Bertram J. Wilm, Klaas P. Pruessmann, S. Johanna Vannesjo

## Abstract

Spiral imaging is very well suited for functional MRI, however its use has been limited by the fact that artifacts caused by gradient imperfections and B_0_ inhomogeneity are more difficult to correct compared to EPI. Effective correction requires accurate knowledge of the traversed k-space trajectory. With the goal of making spiral fMRI more accessible, we have evaluated image reconstruction using trajectories predicted by the gradient impulse response function (GIRF), which can be determined in a one-time calibration step.

GIRF-predicted reconstruction was tested for high-resolution (0.8 mm) fMRI at 7T. Image quality and functional results of the reconstructions using GIRF-prediction were compared to reconstructions using the delay-corrected nominal trajectory and concurrent field monitoring.

The reconstructions using nominal spiral trajectories contain substantial artifacts and the activation maps contain misplaced activation. Image artifacts are substantially reduced when using the GIRF-predicted reconstruction, and the activation maps for the GIRF-predicted and monitored reconstructions largely overlap. The GIRF reconstruction provides a large increase in the spatial specificity of the activation compared to the nominal reconstruction.

The GIRF-reconstruction generates image quality and fMRI results similar to using a concurrently monitored trajectory. The presented approach does not prolong or complicate the fMRI acquisition. Using GIRF-predicted trajectories has the potential to enable high-quality spiral fMRI in situations where concurrent trajectory monitoring is not available.

**Highlights:**

- This work investigates the feasibility of using a one-time system calibration to account for k-space trajectory deviations in spiral fMRI.
- This versatile calibration is based on a linear time-invariant gradient model, the gradient impulse response function (GIRF).
- We show that the image quality and the spatial specificity of the fMRI activation are substantially improved when using the GIRF-predicted trajectories.
- Basing reconstructions on nominal gradient inputs, on the other hand, induces image artifacts and misplaced fMRI activation.
- We demonstrate that system characterization via the GIRF can enable spiral fMRI in situations where concurrent trajectory monitoring is unavailable.

## Introduction

Blood Oxygen Level Dependent (BOLD) functional magnetic resonance imaging (fMRI) requires fast imaging, for which acquisitions with Echo Planar Imaging (EPI) readouts are currently used as the gold standard. Spiral readouts (Ahn et al., 1986) have many desirable properties for rapid acquisitions and have long been considered a promising alternative to EPI for fMRI (Glover, 2012): They can provide higher k-space sampling efficiency compared to EPI sampling (Glover, 2012; Glover and Lee, 1995; Noll et al., 1995), translating into higher resolution within a given readout time, and they also allow for a more flexible choice of echo time (TE). The combination is especially useful for high-resolution fMRI at 7T or above, where for EPI large parallel imaging factors or Partial Fourier are required to achieve the optimal TE for BOLD contrast. Further, spiral imaging has naturally reduced sensitivity to pulsatile motion (Glover and Lee, 1995; Yang et al., 1998), and spiral-in/out trajectories (Glover and Law, 2001) can improve fMRI in regions prone to dropout, such as the orbitofrontal cortex. Finally, spiral sampling is more amenable to high undersampling factors, as the point spread function results in relatively incoherent aliasing (Wright et al., 2014), which can be less detrimental to image quality compared to coherent aliasing, which occurs in undersampled EPI.

Despite these advantages spiral imaging has not yet become a mainstream fMRI acquisition strategy. The reasons for the slow uptake of spiral fMRI include the fact that artifacts caused by gradient imperfections (discrepancy between the actual and nominal gradients) and B_0_ inhomogeneities are more difficult to correct for spiral trajectories compared to EPI. Localized off-resonance resulting from susceptibility-induced field inhomogeneities cause dropout and shifts in EPI, for which a number of established correction methods exist (Andersson et al., 2003; Smith et al., 2004). Similarly for gradient infidelity, the correction of the Nyquist ghost artifact in EPI is considered a routine step in image reconstruction, for example using navigator lines acquired before the readout (Schmitt et al., 1998). For spiral readouts, B_0_ inhomogeneities and gradient imperfections cause blurring, geometric distortions and dropout. To improve gradient fidelity, it is common to perform a delay correction in spiral imaging (Bhavsar et al., 2014; Börnert et al., 1999; Robison et al., 2010). However this typically requires extra calibration scans, and is usually not as effective as the EPI delay correction. Moreover, unlike EPI, the effects of B_0_ inhomogeneities are difficult to correct via image processing methods, and are therefore commonly not addressed at all. As a result, spiral fMRI images have often been blurry, especially around the air-tissue interfaces in the frontal sinuses and ear canals.

Monitoring of the encoding fields during the acquisition using Nuclear Magnetic Resonance (NMR) field probes (Barmet et al., 2008; De Zanche et al., 2008) allows precise measurement of the traversed k-space trajectory. In conjunction with a B_0_ field map covering the imaging FOV, included in a model-based reconstruction, this has been shown to yield high-quality spiral imaging (Engel et al., 2018; Kasper et al., 2018; Wilm et al., 2016), enabling high-resolution spiral fMRI (Kasper et al., 2019). However, the concurrent monitoring requires an additional hardware setup, which is not always available. One alternative to direct field measurements is to model the behavior of the gradient chain. With the appropriate model, deviations from the prescribed encoding that are reproducible (for example, induced by eddy currents) can be measured and corrected for. It has previously been shown (Addy et al., 2012; Vannesjo et al., 2013) that the gradient chain can be considered as a linear, time invariant (LTI) system to a high degree of accuracy. For an LTI system, the relation between the input to the system and its output is determined by the impulse response of the system – in the case of the gradient chain, the gradient impulse response function (GIRF). The GIRF of a specific system can be characterized in a one-time calibration procedure, which then enables to predict the actual gradient output to arbitrary input pulses. The GIRF-predicted output can be used as basis for image reconstruction, which has been shown to yield high quality images for a range of different trajectories (Addy et al., 2012; Campbell-Washburn et al., 2016; Vannesjo et al., 2016).

We have previously demonstrated that GIRF-prediction enables single-shot spiral images with only minor quality differences to using the monitored trajectory (Vannesjo et al., 2016). This evaluation was performed on individual images. In fMRI, however, we perform high duty cycle imaging over extended periods of time (5-10 minutes for a typical fMRI run with a single fMRI session often containing multiple runs). We know that there are long-term effects, for example gradient heating, that violate the LTI assumption at the basis of the GIRF prediction. But we do not know to what extent this will affect an image time-series, such as required for fMRI. We also do not know how the resulting imperfections propagate into the fMRI analysis. The aim of the present work is to evaluate the utility of GIRF-based reconstruction for spiral functional MRI. The results are assessed by comparison with reconstructions based on concurrent field monitoring and nominal trajectories with gradient delay correction.

## Methods

All data were acquired on a 7T Achieva system (Philips Healthcare, Best, Netherlands) using a quadrature-transmit coil and 32-channel head receive array (Nova Medical, Wilmington, MA). The manufacturer’s built-in eddy current compensation was kept activated for all experiments.

### LTI gradient model

A linear and time-invariant system can be described via its impulse response function, which is the output of the system to a very brief input pulse. Knowledge of the system’s impulse response allows predicting the system response *o(t)* to any input, via convolution of the input waveform *i(t)* with the impulse response (equation [1]). In the frequency domain this corresponds to a multiplication with the Fourier transform of the impulse response [2] (typically called the system transfer function – for simplicity we use the acronym *GIRF* referring to the gradient response function both in the time and in the frequency domain).

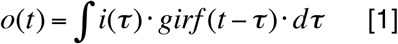

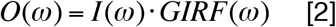

The characterization of the gradient chain was performed similarly as described by Vannesjo et al. (Vannesjo et al., 2013): A set of gradient input pulses were played out and the resulting magnetic fields were measured with a dynamic field camera (Dietrich et al., 2016a) consisting of 16 ^1^H NMR field probes distributed on the surface of a sphere of 10 cm radius. This allows fitting of spherical harmonic basis functions up to 3^rd^ order to the probe measurements. The GIRF was calculated via frequency-domain division of the measured output by the known inputs, using least-squares combination of data from different input pulses. For an accurate GIRF calibration the input gradient pulses should cover the entire range of expected frequencies, while complying with hardware and acquisition time constraints. This was achieved by using 12 different triangular pulses (slew rate 200 T/m/s, time-to-peak 20–158 ms at ~12-ms increments). The GIRF measurements took approximately 3 minutes (12 gradient pulses, 3 gradient directions, 4 averages, 1.2 s TR). The individual probe signals were corrected for concomitant fields terms which are a known deviation from the LTI assumption (for details see (Vannesjo et al., 2016)). This correction was also applied to the concurrent field monitoring data described in the next section.

The LTI model of the gradient system was subsequently used to estimate actual gradient time courses of the imaging acquisitions through a frequency-domain multiplication of the nominal gradient time courses with the measured GIRFs.

### Concurrent field monitoring

Concurrent field monitoring was performed during all imaging acquisitions using ^19^F NMR field probes positioned between the transmit coil and the 32-channel receive array (as shown in (Engel et al., 2018), Fig.1). The probe data was fitted to up to 1^st^ order spherical harmonics producing linear gradient field terms in the three orthogonal directions (and the corresponding k-space trajectory k_x_,k_y_,k_z_), as well as a 0^th^-order field term (and the corresponding phase term, k_0_), which reflects global field changes over time.

The field probes were excited before the start of the readout gradient and the probe signal was acquired concurrently with the imaging readout. Due to the long readout and the strong imaging gradients, the probe signal can de-phase prior to the end of the monitoring period. Each probe’s signal was therefore visually inspected and if the probe signal had very low amplitude and the signal phase exhibited discontinuities the probe was excluded from the spherical harmonic fit. Per subject, between 5-7 probes (on average 6.2) were excluded in this study, leaving approximately 10 probes for the fit.

**Figure 1:**
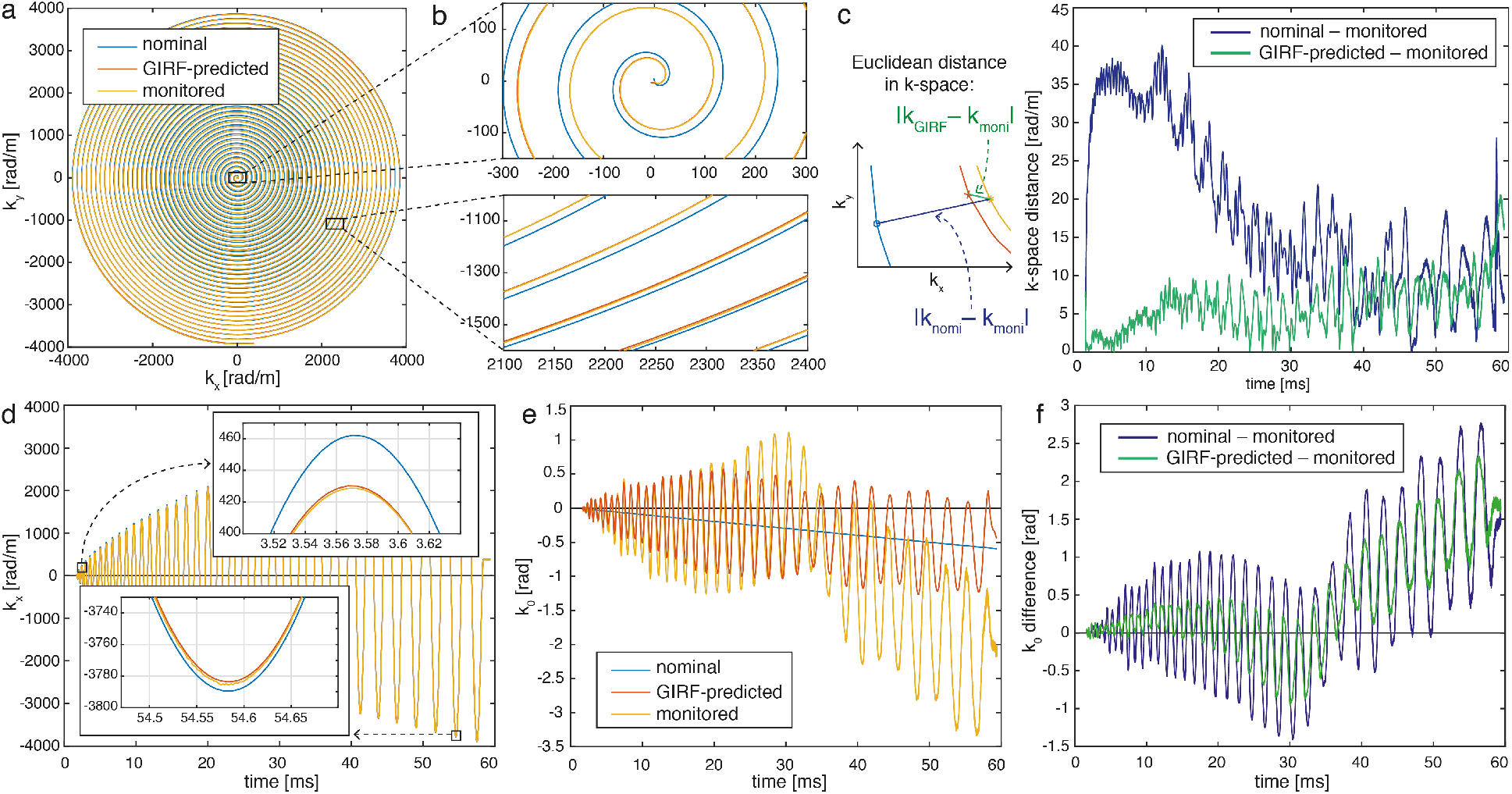
Comparison of a nominal delay-corrected, GIRF-predicted and concurrently monitored spiral trajectory for subject 4 (first slice of first volume). (a) An example spiral trajectory and (b) zooms highlighting differences between the three trajectories. (c) The k-space distance between nominal/GIRF-predicted and monitored trajectories is shown to quantify differences over the course of one spiral readout. (d) k_x_ during spiral readout including zooms. (e) The zeroth order field term, k_0_, over one readout. Note that the nominal k_0_ has a non-zero slope due to the retrospective “frequency adjustment” correction applied. (f) Difference of nominal/GIRF-predicted to measured k_0_.

Undesired saturation of the NMR field probe signal can occur if the repetition time of the probe excitation is short relative to the T_1_ of the probes. To allow sufficient time for signal recovery between measurements, the field camera recording was performed on every third slice. For the non-monitored slices the k-space trajectory from the last monitored slice was used.

### fMRI acquisition

The raw coil data and concurrently monitored trajectories analyzed in this work were acquired as part of a recent study exploring the use of concurrently monitored single-shot spirals for fMRI (Kasper et al., 2019). The dataset contains acquisitions from seven healthy volunteers who had giving written informed consent and were scanned with approval of the local ethics committee. The visual fMRI paradigm used a simple retinotopic mapping protocol (Warnking et al., 2002). It was designed to stimulate quarter-fields of the visual cortex, similar to the one used in (Kasper et al., 2014). The subjects were presented with 15 s blocks of two flickering, black-and-white checkerboard-filled 90° wedges separated by 180°, interleaved with 15 s of rest (fixation cross). Alternating blocks of upper left/lower right (ULLR) and blocks of upper right/lower left (URLL) wedges were presented over 100 volumes (~330s). The subjects were instructed to fixate on a point at the center between the wedges. In order to maintain the subjects’ attention, they were asked to respond to any contrast alteration of the fixation point via a button box.

Images were acquired with a multi-slice 2D gradient-echo sequence with a single-shot Archimedean spiral-out readout (designed according to (Lustig et al., 2008)) of 57 ms duration. The radial spacing of samples was chosen to undersample k-space by a factor of 4 with respect to the field-of-view (FOV) of 23 cm. The transversal images were acquired with an in-plane resolution of 0.8 mm isotropic and a TE of 20 ms, selected for good BOLD contrast. 36 slices of 0.89mm thickness (with a slice gap of 0.11 mm) were acquired, resulting in a FOV of 23×23×3.6 cm and a volume TR of 3.3 s. Excitations were preceded by a Spectral Presaturation with Inversion Recovery (SPIR) fat suppression module (Kaldoudi et al., 1993).

A Cartesian multi-echo GRE scan (FOV = 23 cm, resolution = 1 mm isotropic, TE_1_=4 ms, ∆TE = 1 ms, 6 echoes) was collected to estimate coil sensitivities and B_0_ maps. The first echo was used to estimate the coil sensitivities, by dividing the single-coil images by the root-sum-of-squares coil combination. The B_0_ maps were calculated by voxel-wise fitting of the signal phase over the different echoes. Both the coil sensitivity maps and the B_0_ maps were spatially smoothed before use in subsequent image reconstructions.

One subject was excluded from further analysis due to reduced signal in multiple channels of the head receive array. The data from the six remaining volunteers were reconstructed and analyzed as described below.

### Image reconstruction and fMRI analysis

The images were reconstructed offline in Matlab (MathWorks, Natick, MA, USA) using CG-SENSE (Pruessmann et al., 2001) with multi-frequency interpolation for fast off-resonance correction (Man et al., 1997; Sutton et al., 2003). The full reconstruction model takes into account coil sensitivities, the static field as well as field dynamics over time (for details see (Engel et al., 2018; Kasper et al., 2019)). For each data set three reconstructions were performed using the following k-space trajectories:

1. Delay-corrected nominal trajectory (labeled *nominal* in figures)
2. GIRF-predicted trajectory (labeled *GIRF* or *GIRF-predicted* in figures)
3. Trajectory measured with concurrent field monitoring (labeled *monitored* in figures).

For the monitored and GIRF-predicted reconstructions the imaging data were demodulated by the measured/predicted 0^th^-order phase terms k_0_. It has previously been demonstrated that demodulation with an accurate estimate of k_0_ can substantially improve image quality (Vannesjo et al., 2016).

A center frequency adjustment is a typical fMRI pre-scan, which was not performed in this study because it is redundant when using concurrent field monitoring. In order to not artificially disadvantage the GIRF-predicted and nominal reconstructions, we performed a processing step equivalent to frequency adjustment. The center frequency was determined once for each fMRI time series via a linear fit on the first 0.9 ms of the monitored k_0_ (using the data from first slice of the first volume, before the readout gradient starts). The imaging data for the entire fMRI time series was then demodulated by this center frequency for the nominal and the GIRF-predicted reconstruction.

The nominal trajectories were delay-corrected prior to reconstruction. We minimized the RMSE trajectory difference between the nominal and the monitored trajectory on the first readout of the time series. A global delay was chosen as the small differences between the gradient axes were within the standard deviation of the delay calibration.

The reconstructed image time series was corrected for subject translations and rotations using MCFLIRT (Jenkinson et al., 2002) in the FMRIB Software Library (FSL) (Jenkinson et al., 2012) and was pre-whitened using FILM/FSL (Woolrich et al., 2001). The GLM (analyzed using FEAT/FSL (Jenkinson et al., 2012)) contained the regressors for the ULLR and the URLL stimulation blocks convolved with a Gamma function. Activation was assessed using z-statistics contrasting ULLR versus URLL ([1 −1] in the design matrix). In order to produce fMRI data with high spatial specificity we performed no spatial smoothing or clustering. We report activation maps at a liberal threshold of z > 2.3 (p < 0.01).

All of the analysis was performed on a per-subject basis in the space of each subject’s functional data to avoid any degradation of the spatial resolution by registration. The first echo of the multi-echo GRE scans was registered to the functional data and used as the subjects’ structural image. For the analysis of the functional results, masks of the grey matter (GM) and white matter (WM) in the visual cortex were determined as an intersection of a V1-V3 mask (using Juelich atlas (Schleicher et al., 2005; Zilles and Amunts, 2010) labels 81-86 in FSL) and subject-specific GM/WM masks generated by segmenting the structural image using FAST/FSL (Zhang et al., 2001). The GM/WM masks were generated by conservatively thresholding the partial volume maps at 0.8 to exclude most partial volume voxels from the masks. In Table 1 the average and 90^th^ percentile of the absolute value of significant (z > 2.3) z-stats within the GM V1-V3 ROI are reported.

**Table 1:** Results summary metrics for all subjects (from top to bottom): RMSE trajectory errors for the k-space distance and the zeroth order field term, k_0_, using the monitored trajectory as the ground truth, RMSE image error using the monitored reconstruction as the reference, average tSNR in the brain, average and 90^th^ percentile of significant z-statistics in the GM V1-V3 ROI and the AUC from the ROC plots to evaluate spatial specificity of the z-statistic maps.

|  | Subjects → | 1 | 2 | 3 | 4 | 5 | 6 | Mean |
| --- | --- | --- | --- | --- | --- | --- | --- | --- |
| Trajectories | RMSE k-space distance nomi [rad/m] | 19.78 | 19.60 | 20.84 | 19.63 | 18.67 | 19.40 | 19.66 |
|  | RMSE k-space distance GIRF [rad/m] | 9.85 | 10.66 | 12.80 | 9.07 | 10.55 | 12.19 | 10.85 |
| | RMSE $k_0$ nomi [rad] | 1.186 | 0.89 | 1.078 | 1.04 | 1.34 | 0.85 | 1.06 |
| | RMSE $k_0$ GIRF [rad] | 1.11 | 0.75 | 0.90 | 0.84 | 1.24 | 0.81 | 0.94 |
| Images | RMSE nomi [%] | 6.34 | 6.45 | 5.22 | 6.97 | 5.03 | 5.06 | 5.85 |
|  | RMSE GIRF [%] | 2.01 | 2.07 | 1.98 | 2.36 | 2.52 | 1.95 | 2.15 |
| tSNR | tSNR moni | 16.44 | 19.39 | 14.98 | 15.43 | 13.95 | 15.4 | 15.93 |
|  | tSNR GIRF | 14.94 | 18.25 | 14.23 | 14.71 | 13.37 | 14.7 | 15.03 |
|  | tSNR nomi | 15.47 | 19.42 | 14.90 | 15.66 | 14.09 | 15.5 | 15.84 |
| fMRI | mean zstat moni | 3.74 | 4.09 | 3.98 | 4.33 | 4.04 | 3.72 | 3.98 |
|  | mean zstat GIRF | 3.66 | 3.92 | 3.86 | 4.14 | 3.73 | 3.75 | 3.84 |
|  | mean zstat nomi | 3.54 | 3.94 | 3.85 | 3.84 | 3.55 | 3.81 | 3.76 |
|  | 90<sup>th</sup> perct. moni | 5.72 | 6.74 | 6.34 | 7.13 | 6.62 | 5.76 | 6.39 |
|  | 90<sup>th</sup> perct. GIRF | 5.58 | 6.35 | 6.04 | 6.74 | 5.77 | 5.88 | 6.06 |
|  | 90<sup>th</sup> perct. nomi | 5.14 | 6.20 | 5.83 | 5.97 | 5.32 | 5.94 | 5.73 |
|  | ROC AUC moni | 928 | 758 | 860 | 1062 | 624 | 755 | 831 |
|  | ROC AUC GIRF | 809 | 615 | 703 | 981 | 577 | 723 | 735 |
|  | ROC AUC nomi | 548 | 341 | 266 | 295 | 80 | 463 | 332 |

The concurrently monitored trajectories and the resulting image reconstructions were used as reference to assess the nominal and GIRF-predicted data. The trajectory error and image artifacts were accordingly quantified as root-mean-squared error (RMSE) compared to concurrent monitoring (Table 1), and the GIRF/nominal z-statistic maps were compared to the ones derived from concurrent monitoring data. Additionally, receiver operator characteristic (ROC) curves were used to provide an assessment of the fMRI results without selecting the monitored reconstruction as the ground truth. ROC analysis typically involves plotting the number of true-positives against the number of false-positive findings. In this work we used the subject-specific gray and white matter masks of V1-V3 to identify “true positive” and “false positive” activation respectively and plotted this while varying the z-statistic threshold from 0 to the maximum z present in the data. The area under the curve (AUC) gives a measure of spatial specificity and was used to compare between the reconstructions. This analysis has two major advantages: i) it provides a quantitative assessment of the activation maps without requiring one reconstruction as ground truth, and ii) it encompasses the full z-statistics instead of relying on a specific significance threshold (for example z > 2.3).

The temporal SNR (tSNR) was evaluated in the motion-corrected fMRI time series, and was calculated on a voxel-by-voxel basis as the mean signal over time divided by the temporal standard deviation of the signal. The tSNR was averaged over the GM V1-V3 ROI and reported in Table 1 for each subject and reconstruction.

### Code and data availability

The data from subject 6 will be made available on ETH Research Collection upon publication. This will include the raw coil data in ISMRMRD format, the nominal, GIRF-predicted and monitored k-space trajectories and the corresponding reconstructed image time series. The raw data sets from the other subjects cannot be made publicly available, as we did not obtain explicit subject consent to share data for these subjects. However, we provide the mean spiral fMRI images for all three reconstructions with the corresponding activation maps for all subjects on NeuroVault (https://identifiers.org/neurovault.collection:6526).

The image reconstruction in this work was performed using custom Matlab implementation of CG-SENSE (Pruessmann et al., 2001) algorithm. A demo version of the reconstruction pipeline is publicly available on GitHub (https://github.com/mrtm-zurich/rrsg-arbitrary-sense), however without the multi-frequency interpolation used for the B_0_ correction in this work.

Scripts (bash and Matlab) for the post-processing and analysis pipeline will be available on https://github.com/MRI-gradient/paper-GIRF-spiral-fMRI. Matlab code for GIRF calculation and trajectory prediction using the GIRF can also be found in the same repository.

## Results

The delay-corrected nominal spiral trajectories deviate substantially from the ones measured with the NMR field probes, especially close to the center of k-space where the gradients are rapidly changing (Fig.1 a-d). For the example subject shown in Fig. 1 the distance between the nominal and the measured trajectories reach ~1.5*1/FOV. The GIRF-predicted spiral trajectories follow the measured ones much more closely, especially during the first 40 ms of the readout. Towards the edge of k-space, however, there is little improvement from the GIRF-predicted trajectories over the nominal ones. For the example subject in Fig. 1 the maximum k-space deviation between the GIRF-predicted and the measured trajectories is ~0.75*1/FOV, with the largest deviations occurring at the end of the readout.

The root-mean-square trajectory error (RMSE), defined here as the Euclidian distance to the monitored trajectory, averaged over all slices and volumes, is reported for all subjects in Table 1. Averaged over all subjects, the nominal RMSE and GIRF-predicted RMSE were 19.66 rad/m and 10.85 rad/m respectively. This corresponds to a ~45% RMSE reduction when using GIRF prediction over nominal trajectories. Figure 1e/f show the concurrently monitored, GIRF-predicted and nominal k_0_, reflecting global temporal variations in B_0_. The measured k_0_ oscillations are closely coupled with those of the spiral readout gradients. This is partially predicted by the GIRF, but the amplitude of oscillations is not captured accurately, especially towards the end of the readout. Additionally, the monitored k_0_ exhibits slower trends over the readout, for example a change in slope at about 35 ms. These slower dynamics are qualitatively similar across subjects but are not captured by the GIRF prediction, suggesting that they are not linearly related to the gradient waveform. Average RMSE over all subjects (see Table 1 for results for individual subjects) is 1.06 rad for nominal k_0_ and 0.94 rad for GIRF-predicted k_0_, which corresponds to an improvement of 11%.

The GIRF-predicted and nominal trajectories are the same for all reconstructions within a time series, whereas the monitored trajectory is updated every 3^rd^ readout. Fig. 2 illustrates how the monitored trajectory changes over the course of the fMRI experiment. Over the 5.5 minute experiment the trajectory gradually shifts from the first volume (blue) to the last volume (red), likely due to gradient heating. Note that the shift, however, is small compared to the distance to the nominal trajectory. The RMSE for the GIRF-predicted vs. monitored trajectories increases over the course of the experiment, but remains considerably below the RMSE of the nominal vs. measured trajectories (Fig. 2b). Interestingly, the latter slightly improves during the experiment. For the measured k_0_ (Fig. 2c/d) the amplitude of the observed oscillations remains fairly consistent buy the slope of the k_0_ drift over the readout changes substantially.

**Figure 2:**
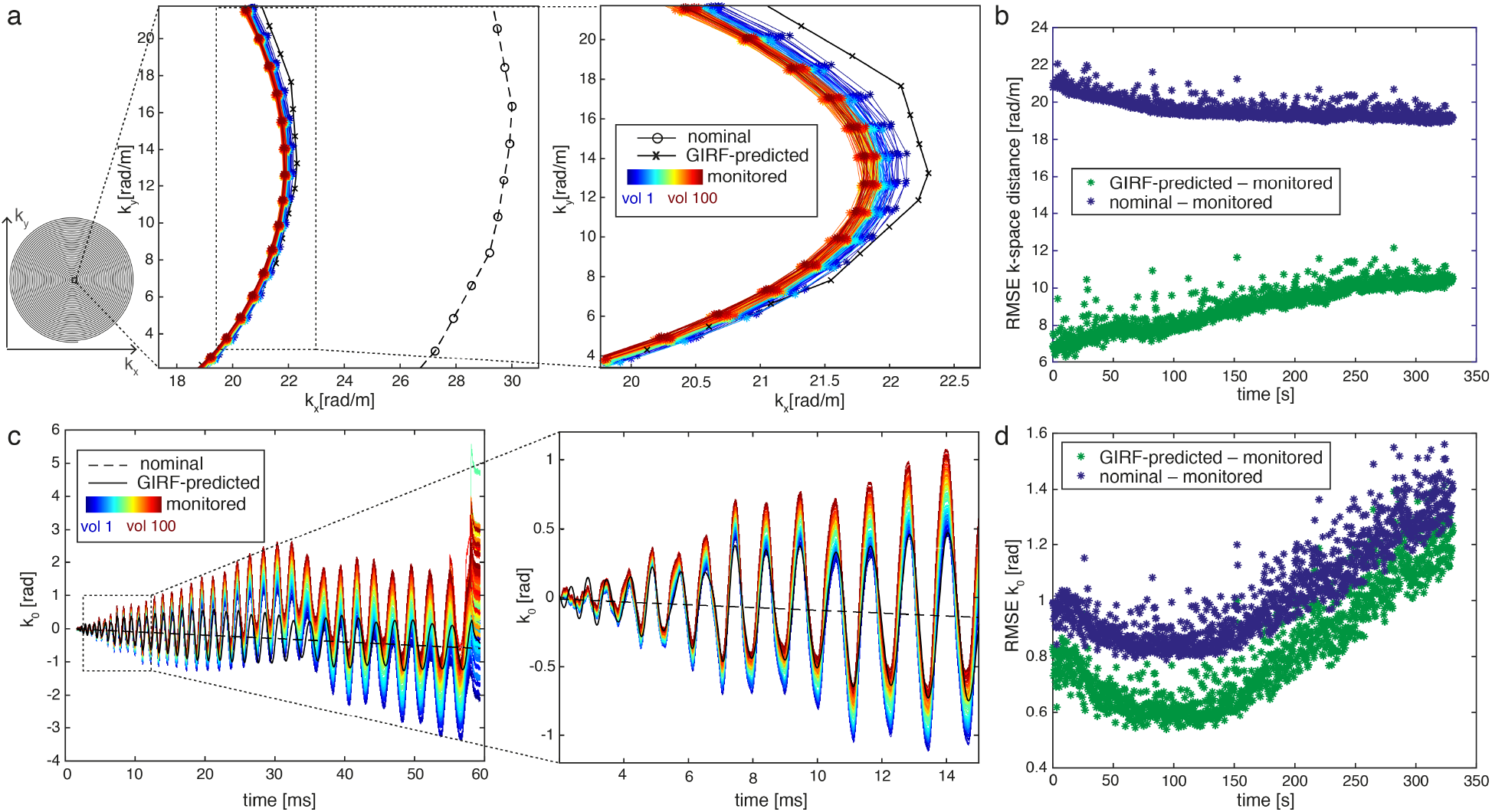
Evolution of k-space trajectories (for subject 4) over the fMRI experiment showing how (a) a monitored spiral trajectory changes from the first volume (dark blue) to the last (dark red) including zoom and (b) RMSE on the k-space distance for the nominal/GIRF-predicted trajectories with respect to the monitored one. In the lower row (c) k_0_ with zoom and (d) RMSE of the nominal/GIRF-predicted k_0_ with respect to the measured one are shown.

The nominal spiral images are heavily corrupted by blurring and geometric distortion (Fig. 3). The GIRF-predicted reconstruction provides much improved image quality. Residual artifacts (mainly subtle blurring and some ringing) can be observed in the difference images to the monitored reconstruction. The global image artifact levels, defined here as the RMSE to the monitored reconstruction averaged over all voxels in a brain mask and all volumes in the fMRI time-series, are reported in Table 1 (image differences reported as percent of the maximum value in the monitored reconstruction). Averaged over all subjects the artifact level was 5.85% for nominal image reconstructions and 2.15% for GIRF-predicted trajectories, which corresponds to an improvement of 63%.

**Figure 3:**
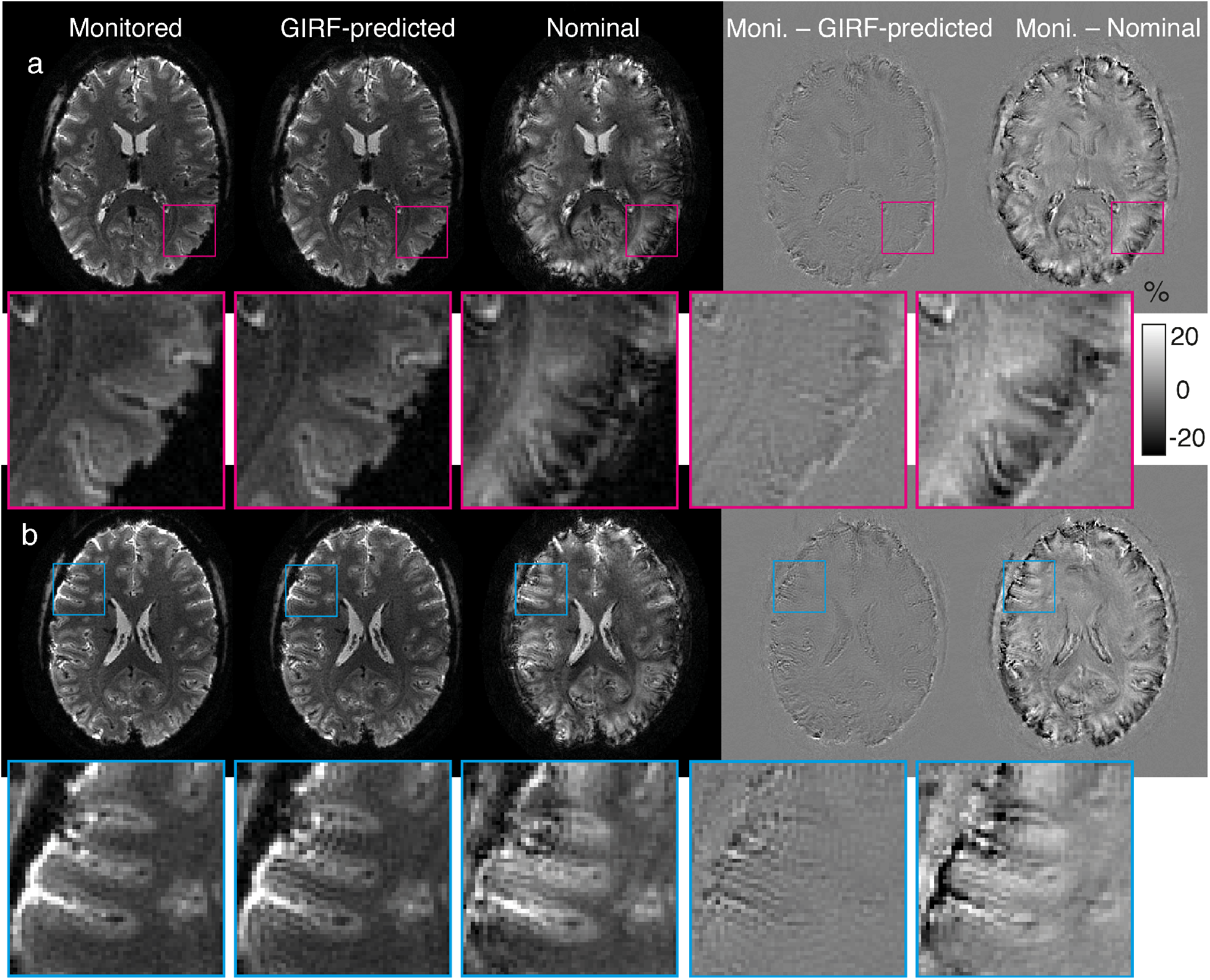
Image quality comparison of reconstructions using monitored trajectories (left), GIRF-predicted trajectories (centre) and nominal trajectories (right) for two different slices for subject 6 (first volume of the time series). To the right of each sub-figure, the differences to the corresponding reconstructions based on the monitored trajectory are displayed. The difference images are scaled to percent of the maximum value in the monitored reconstruction. In the inferior slice (a) the GIRF-predicted reconstruction only contains a small increase in blurring compared to the monitored reconstruction. In a superior example slice (b) the GIRF-predicted reconstruction additionally exhibits an increase in ringing artifacts.

A comparison of the first and final volume in the fMRI time series revealed an increase in image artifacts over time for the GIRF reconstruction (shown for an example subject in Fig. 4), in line with the observed increase in trajectory error. The image quality however remained much improved over the nominal spiral images through-out the time series, with an image artifact reduction compared to nominal of 65% in the first volume and 57% in the final volume (averaged over all subjects).

**Figure 4:**
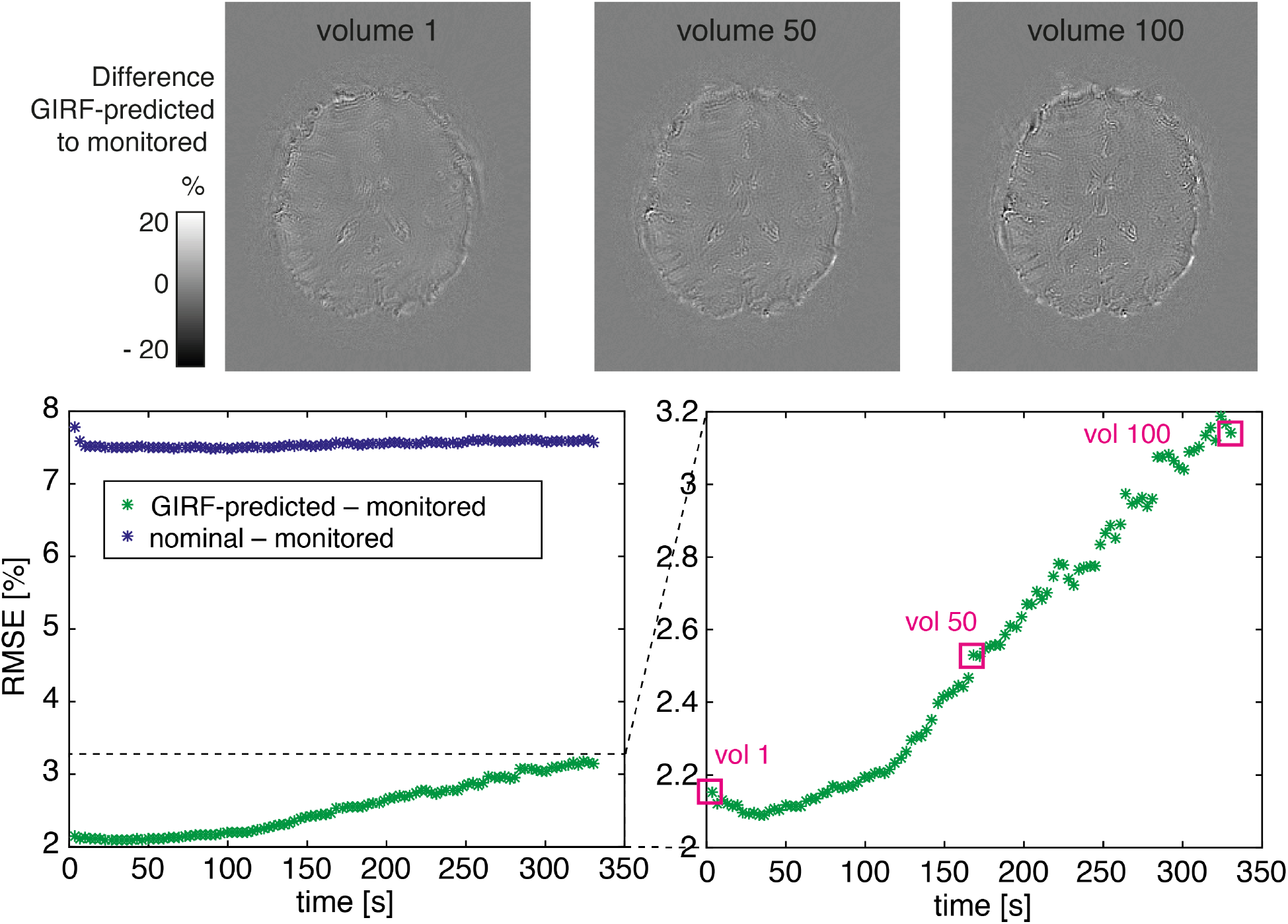
Quality of GIRF-predicted reconstructions over the fMRI time-series (shown for subject 4): Top row shows difference images of monitored and GIRF-predicted reconstructions at the start, middle and the end of the 330-second fMRI acquisition. The bottom row shows the RMSE for GIRF-predicted/nominal reconstructions with respect to the monitored one. A zoom showing the dynamics of the GIRF-predicted RMSE is shown on the bottom right.

The average temporal SNR over all subjects (Table 1) is highest for monitored reconstructions, slightly lower for the nominal reconstructions (<1% reduction) and lowest for the reconstructions using GIRF-predicted trajectories (~5% reduction). The improved tSNR with concurrent monitoring is expected, as we are reducing the temporal variance by correcting for field effects caused by gradient heating and physiological field fluctuations (Bollmann et al., 2017; Bright and Murphy, 2017). This is not captured by the GIRF where we use the same trajectory for each volume. The higher tSNR of the nominal reconstruction as compared to the GIRF reconstruction could potentially be explained by the larger amount of blurring present, which causes local averaging of signal in the image.

Figures 5 and 6 show fMRI activation maps, in a single subject in various orientations, and in a single slice in all subjects, respectively. For the monitored and GIRF-predicted reconstructions, the spiral fMRI results show good correspondence of the activation with gray matter architecture, while the nominal data contain misplaced activation (apparent for example where activation is crossing white matter boundaries). The activation for the GIRF-predicted and monitored reconstructions largely overlap in all subjects, whereas there are substantial deviations in the nominal activation maps, as demonstrated by activation difference maps (Fig. 6).

**Figure 5:**
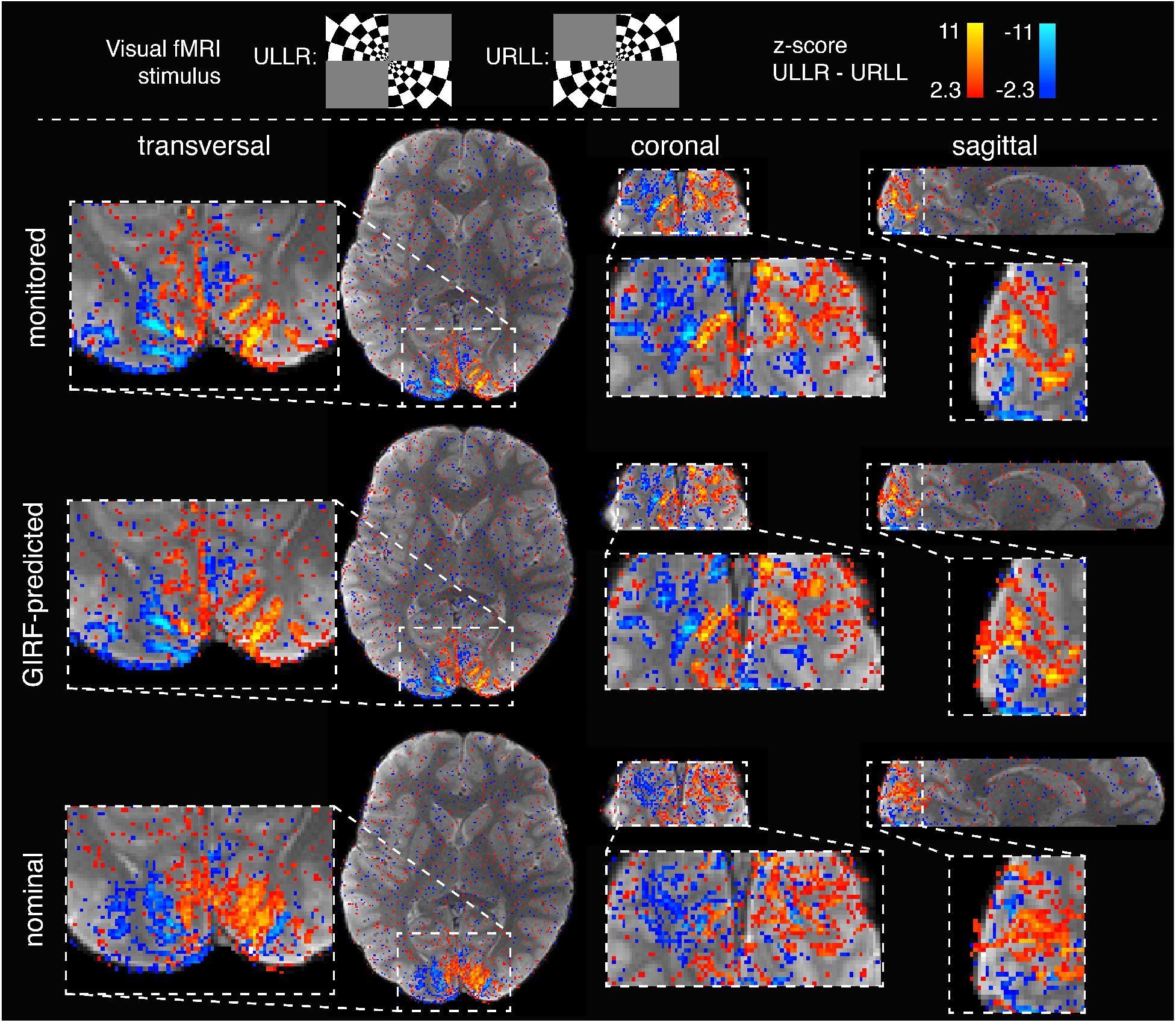
Evaluation of the fMRI experiment designed to stimulate the quarter-fields of the visual cortex. Z-statistic maps (contrasting ULLR versus URLL) overlaid on the structural image (shown for subject 4). The activation for the monitored and GIRF-predicted reconstructions match the grey matter architecture well, as seen for example along the calcarine sulcus (sagittal view), while the nominal reconstruction results in misplaced activation.

**Figure 6:**
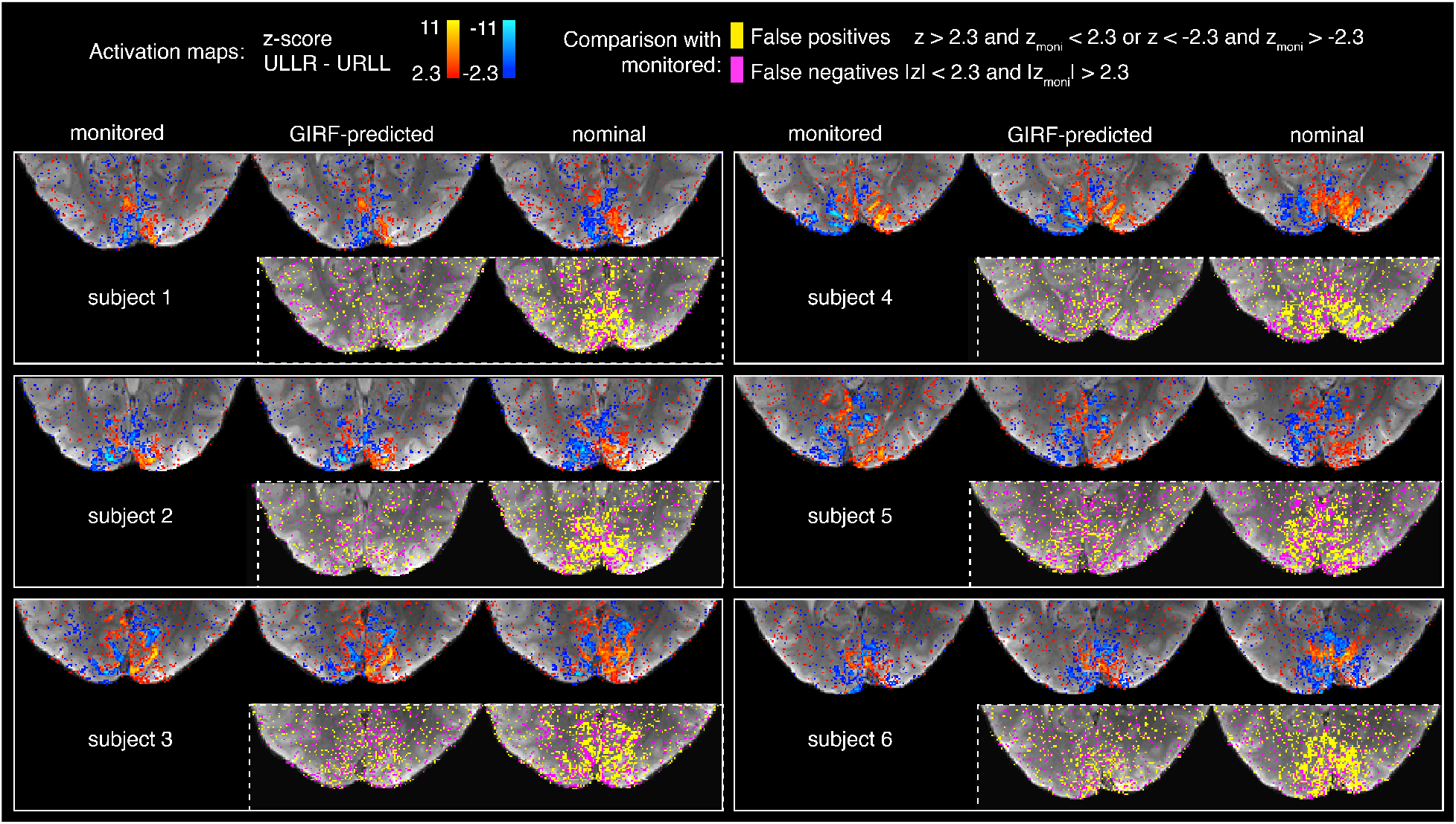
Activation maps (transversal section of visual cortex) for six subjects. Maps of false positives (yellow) and false negatives (pink) with respect to the monitored reconstruction are displayed below each GIRF-predicted and nominal image.

The ROC analysis confirms that the GIRF-predicted reconstructions yield nearly as good spatial specificity as the monitored reconstructions, and considerably better than nominal reconstructions (Fig. 7). Compared to the nominal reconstructions, the GIRF reconstructions yield ~122% increase in specificity as captured by the AUC, averaged over all subjects. The monitored reconstructions in turn provide a smaller additional improvement over the GIRF-reconstructions, with an average increase of the AUC by ~13%. The ROC analysis assumes that all activated voxels within the GM V1-V3 ROI are true positives. Note that this is different to how true positives are defined in the z-statistics map comparison with monitored as the ground truth in Figure 6.

**Figure 7:**
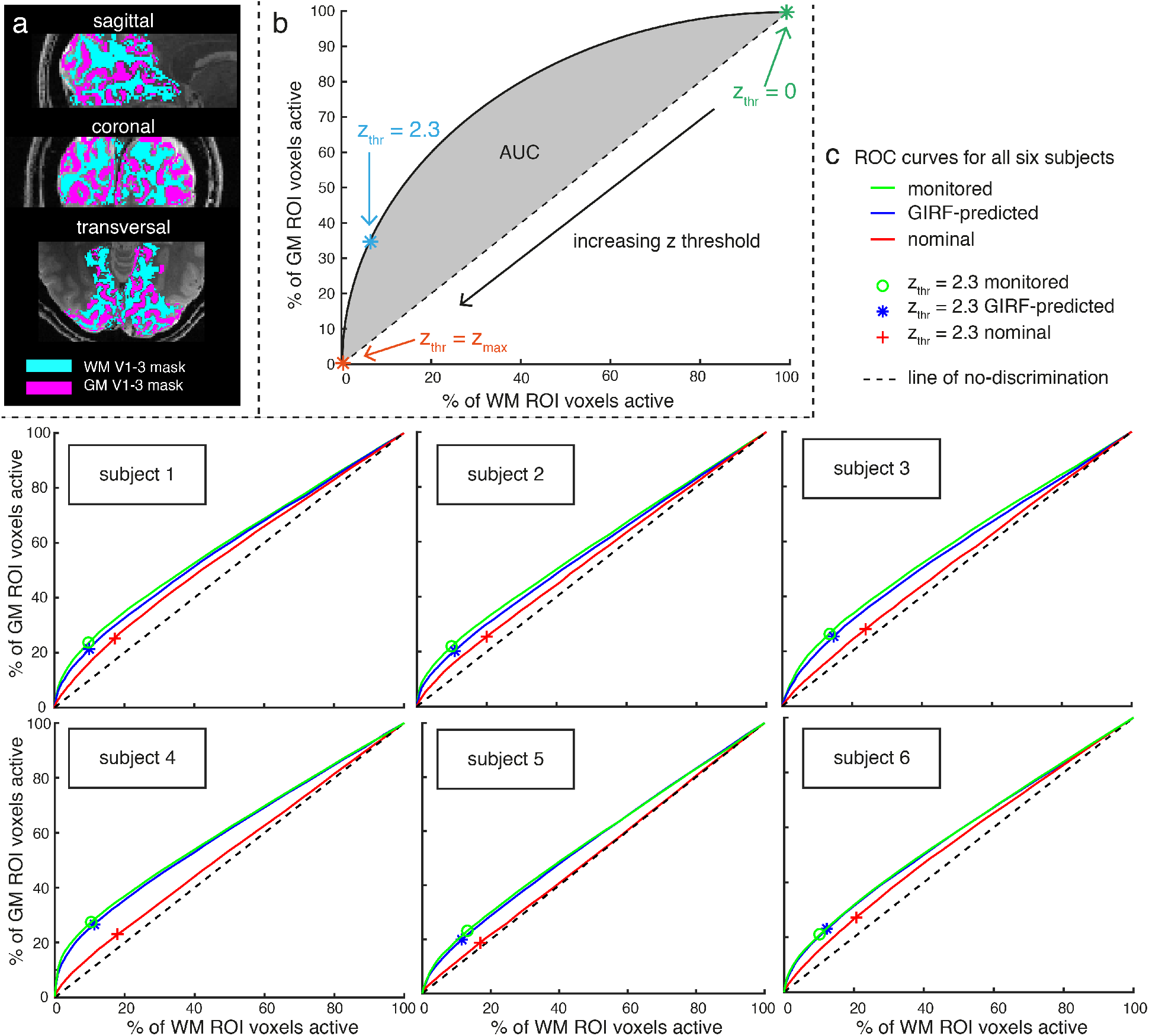
(a) Grey and white matter masks of V1-V3 used for the analysis. (b) Schematic explaining receiver operating characteristic (ROC) curve analysis used to assess the spatial specificity of the different reconstructions (without having to choose a specific one as ground truth). The dashed line is the line of no discrimination, indicating equal amounts of true and false positives. The area under the curves (AUC) values were used as a summary metric and are reported in Table 1. (c) ROC curves for all six subjects, indicating that the spatial specificity of the activation is highest for the monitored reconstructions. The GIRF-predicted reconstructions result in only slightly reduced specificity whereas the nominal curve lies substantially closer to the line of no discrimination.

## Discussion

The main goal of this work was to determine if image reconstruction based on an LTI gradient system model is suitable for use in functional MRI with spiral trajectories. The results presented here first confirmed the conclusion of previous work, showing that reconstructions using the nominal trajectory contain a large amount of artifacts (blurring, ringing and distortion) while monitored and GIRF-predicted trajectories yield high-quality images. The image quality of the GIRF-predicted reconstructions degraded slightly over the course of the time series, but remained superior to reconstructions based on nominal trajectories. The higher image quality translated into increased spatial specificity in the fMRI analysis, yielding activation patterns that closely followed the gray/white matter architecture in the visual cortex. Nominal reconstructions, on the other hand, yielded misplaced activation that was not localized to the gray matter, due to the blurring and other artifacts present in the images.

The long high-duty cycle acquisitions in fMRI cause heating of the gradient coils and surrounding structures. For our acquisition we observed gradient temperature increases of ~10-15 °C over a single ~5 minute fMRI run and ~25-30 °C increase over the entire scanning session consisting of four fMRI runs and a number of shorter scans. Such a substantial temperature increase alters electrical and mechanical material properties and thus changes the behavior of the gradient chain. This is a deviation from the LTI assumption underlying the GIRF approach, which could explain the observed increase in artifacts in the GIRF-predicted reconstructions over the course of the fMRI acquisition. The GIRF measurement is relatively low duty cycle and was performed starting from a cold state of the system. Therefore, towards the end of an fMRI time series the gradients are in a different thermal state to the one they were characterized in. Overall the impact of this on the fMRI results was small. The spatial specificity of the GIRF-predicted fMRI results was very close to the monitored ones, while it was substantially reduced for the nominal reconstruction. In future work, the GIRF model may be further improved by incorporating temperature-dependent GIRFs (Dietrich et al., 2016b; Nussbaum et al., 2018; Stich et al., 2019). The hardware temperature can easily be assessed via the scanner’s temperature monitoring system or using separate temperature sensors and this information can then be used to select the optimal GIRF for each measurement. Alternatively the GIRF approach could be combined with additional navigators that, for example, track global field changes per slice or volume. This would allow adjusting the slope of k_0_, which we had observed to vary substantially over the course of the fMRI time series (Fig. 2).

The delay correction of the nominal trajectory did not improve the image quality much, as compared to no delay correction (data not shown). This stands in contrast to EPI where calibrating a delay between odd and even lines typically allows substantial artifact reduction. The delay of the gradient chain is dependent on the frequency of the input waveform, and is therefore not a single parameter valid for all gradient waveforms (Vannesjo et al., 2013). The spiral readout gradients sweep a large range of temporal frequencies, whereas the EPI has a dominant peak at the switching frequency. Presumably for this reason, a single delay correction works well in EPI, whereas for spiral imaging it is important to know the full response over a large range of frequencies.

B_0_ related artifacts, such as dropout and blurring, scale with field strength, therefore it is especially important to include B_0_ correction for fMRI at 7T. We observed that the artifacts were worst in the nominal reconstructions near air-tissue interfaces, where B_0_ inhomogeneity is large, despite the fact that static B_0_ was accounted for in all reconstructions. This is because B_0_ correction relies on geometric congruency between the field map and the encoded image (Spirig et al., 2017). Accurate knowledge of the encoding fields therefore becomes even more important at ultra-high field.

We evaluated the spatial specificity of the GIRF-predicted reconstruction, to test suitability of the approach for high-resolution fMRI, as for example required to detect activation on the level of cortical laminae (Huber et al., 2017; Kok et al., 2016) and columns (Cheng et al., 2001; Yacoub et al., 2008). We assessed the spatial specificity using a ROC-style analysis, which allows quantifying specificity without choosing one reconstruction as a ground truth and is independent of a specific significance threshold. The challenge with this method is that it requires accurate true/false positive masks, which relies on accurate segmentation and registration of an anatomical atlas to the subjects’ functional data. To obtain trust-worthy masks, we visually inspected the registration and segmentation in each subject, and chose a conservative threshold for the automatic gray/white matter segmentation. Voxels straddling the border between gray and white matter were therefore not included in either of the masks.

The GIRF-predicted trajectories consistently provided good results for all subjects acquired. There were some inter-subject differences in how closely the GIRF-predicted reconstructions matched the monitored ones (see Fig. 5 and 7). The fMRI scans used in this study were acquired at different time points within the scanning session (sometimes it was the first longer scan of the session whereas other times a few other fMRI runs had been performed immediately beforehand). This could potentially explain some of the differences in performance of the GIRF-prediction between subjects.

The reconstructions using measured trajectories provided the best results, both in terms of image quality and fMRI activation patterns. Concurrent monitoring allows capturing dynamic field effects that violate the LTI assumption, including non-linear and time-dependent responses of the gradient system, as well as non-reproducible effects (e.g. caused by the subject). However, concurrent monitoring is technically challenging and can be difficult to incorporate into routine fMRI scanning. The approach relies on an external hardware setup, optimized for the specific purpose of monitoring long readouts at high resolution. This requires field probes with suitable specifications (e.g. size and doping of the probe) to avoid probe signal de-phasing during the measurement. GIRF-prediction provides an alternative when concurrent monitoring is not feasible, for example if the optimal field monitoring setup is not available or if de-phasing still occurs (e.g., due to a poor shim). Moreover, concurrent monitoring could be combined with the GIRF model, where parts of the readout is measured and the rest is filled in by GIRF prediction (Wilm et al., 2019).

GIRF characterization is a one-time calibration step (previous work has shown the GIRF to be stable over at least 3 years (Vannesjo et al., 2016)) and can be performed without any specialized equipment. In this study the GIRF was determined using a dynamic field camera, which allows very accurate characterization of the encoding fields with high frequency resolution, including spatial cross-terms and higher-order terms. However the GIRF can also be measured using a phantom-based approach (Addy et al., 2012; Duyn et al., 1998; Rahmer et al., 2019), at the cost of some loss in the frequency resolution of the GIRF (Graedel et al., 2017a).

We used a spiral-out fMRI protocol for this study, but the GIRF-based trajectory prediction is a generic method that can be used for any trajectory. For example it could be employed for hybrid spiral-in/out methods (Glover and Law, 2001), which can provide high BOLD sensitivity as well as improved signal dropout artifacts. Beyond spirals the GIRF could enable other non-Cartesian fMRI techniques, which require accurate gradient correction, such as radial (Lee et al., 2010), PROPELLER (Krämer et al., 2012) and TURBINE fMRI (Graedel et al., 2017b). Furthermore, the approach presented here may also be useful to correct EPI trajectories, as the GIRF prediction captures effects that the commonly used odd-even lines EPI Nyquist ghost correction schemes (Schmitt et al., 1998) do not address (Vannesjo et al., 2016).

## Conclusion

GIRF-predicted trajectories have the potential to enable high-quality spiral fMRI in situations where concurrent monitoring is not available. The presented approach requires only a one-time calibration per system, thus the fMRI acquisition is not prolonged or complicated by the acquisition of additional data for correction purposes.

## Acknowledgments

We would like to thank the Oxford-Brain@McGill-ZNZ Partnership in the Neurosciences for funding this project (OMZPN/2015/1/3). This work was also supported by the European Union’s Horizon 2020 research and innovation programme under the Marie Sklodowska‐Curie grant agreement No 659263 (J.V.) and the NCCR “Neural Plasticity and Repair” (L.K.). The Wellcome Centre for Integrative Neuroimaging is supported by core funding from the Wellcome Trust (203139/Z/16/Z). Technical support from Philips Healthcare is gratefully acknowledged.

## References

Addy, N.O., Wu, H.H., Nishimura, D.G., 2012. Simple method for MR gradient system characterization and k‐space trajectory estimation. Magn. Reson. Med. 68, 120–129.

Ahn, C.B., Kim, J.H., Cho, Z.H., 1986. High-Speed Spiral-Scan Echo Planar NMR Imaging-I. IEEE Trans. Med. Imaging. https://doi.org/10.1109/TMI.1986.4307732

Andersson, J.L.R., Skare, S., Ashburner, J., 2003. How to correct susceptibility distortions in spin-echo echo-planar images: application to diffusion tensor imaging. Neuroimage 20, 870–888.

Barmet, C., De Zanche, N., Pruessmann, K.P., 2008. Spatiotemporal magnetic field monitoring for MR. Magn. Reson. Med. 60, 187–197.

Bhavsar, P.S., Zwart, N.R., Pipe, J.G., 2014. Fast, variable system delay correction for spiral MRI. Magn. Reson. Med. https://doi.org/10.1002/mrm.24730

Bollmann, S., Kasper, L., Vannesjo, S.J., Diaconescu, A.O., Dietrich, B.E., Gross, S., Stephan, K.E., Pruessmann, K.P., 2017. Analysis and correction of field fluctuations in fMRI data using field monitoring. Neuroimage. https://doi.org/10.1016/j.neuroimage.2017.01.014

Börnert, P., Schomberg, H., Aldefeld, B., Groen, J., 1999. Improvements in spiral MR imaging. Magn. Reson. Mater. Physics, Biol. Med. 9, 29–41.

Bright, M.G., Murphy, K., 2017. Cleaning up the fMRI time series: Mitigating noise with advanced acquisition and correction strategies. Neuroimage. https://doi.org/10.1016/j.neuroimage.2017.03.056

Campbell-Washburn, A.E., Xue, H., Lederman, R.J., Faranesh, A.Z., Hansen, M.S., 2016. Real-time distortion correction of spiral and echo planar images using the gradient system impulse response function. Magn. Reson. Med. https://doi.org/10.1002/mrm.25788

Cheng, K., Waggoner, R.A., Tanaka, K., 2001. Human ocular dominance columns as revealed by high-field functional magnetic resonance imaging. Neuron. https://doi.org/10.1016/S0896-6273(01)00477-9

De Zanche, N., Barmet, C., Nordmeyer-Massner, J.A., Pruessmann, K.P., 2008. NMR probes for measuring magnetic fields and field dynamics in MR systems. Magn. Reson. Med. 60, 176–186.

Dietrich, B.E., Brunner, D.O., Wilm, B.J., Barmet, C., Gross, S., Kasper, L., Haeberlin, M., Schmid, T., Vannesjo, S.J., Pruessmann, K.P., 2016a. A field camera for MR sequence monitoring and system analysis. Magn. Reson. Med. 75, 1831–1840.

Dietrich, B.E., Reber, J., Brunner, D.O., Wilm, B.J., Pruessmann, K.P., 2016b. Analysis and prediction of gradient response functions under thermal load, in: Proc. Intl. Soc. Mag. Reson. Med, Singapore. p. 3551.

Duyn, J.H., Yang, Y., Frank, J.A., van der Veen, J.W., 1998. Simple correction method for k-space trajectory deviations in MRI. J. Magn. Reson. 132, 150–153.

Engel, M., Kasper, L., Barmet, C., Schmid, T., Vionnet, L., Wilm, B., Pruessmann, K.P., 2018. Single-shot spiral imaging at 7 T. Magn. Reson. Med. 80, 1836–1846.

Glover, G.H., 2012. Spiral imaging in fMRI. Neuroimage 62, 706–712.

Glover, G.H., Law, C.S., 2001. Spiral‐in/out BOLD fMRI for increased SNR and reduced susceptibility artifacts 46, 515–522.

Glover, G.H., Lee, A.T., 1995. Motion artifacts in fMRI: comparison of 2DFT with PR and spiral scan methods. Magn. Reson. Med. 33, 624–635.

Graedel, N.N., Hurley, S.A., Clare, S., Miller, K.L., Pruessmann, K.P., Vannesjo, S.J., 2017a. Comparison of gradient impulse response functions measured with a dynamic field camera and a phantom-based technique., in: Proceedings of the ESMRMB. p. 378.

Graedel, N.N., McNab, J.A., Chiew, M., Miller, K.L., 2017b. Motion correction for functional MRI with three-dimensional hybrid radial-Cartesian EPI. Magn. Reson. Med. 78, 527–540. https://doi.org/10.1002/mrm.26390

Huber, L., Handwerker, D.A., Jangraw, D.C., Chen, G., Hall, A., Stüber, C., Gonzalez-Castillo, J., Ivanov, D., Marrett, S., Guidi, M., Goense, J., Poser, B.A., Bandettini, P.A., 2017. High-Resolution CBV-fMRI Allows Mapping of Laminar Activity and Connectivity of Cortical Input and Output in Human M1. Neuron. https://doi.org/10.1016/j.neuron.2017.11.005

Jenkinson, M., Bannister, P., Brady, M., Smith, S., 2002. Improved Optimization for the Robust and Accurate Linear Registration and Motion Correction of Brain Images. Neuroimage 17, 825–841.

Jenkinson, M., Beckmann, C.F., Behrens, T.E.J., Woolrich, M.W., Smith, S.M., 2012. FSL. Neuroimage 62, 782–790.

Kaldoudi, E., Williams, S.C.R., Barker, G.J., Tofts, P.S., 1993. A chemical shift selective inversion recovery sequence for fat-suppressed MRI: Theory and experimental validation. Magn. Reson. Imaging. https://doi.org/10.1016/0730-725X(93)90067-N

Kasper, L., Engel, M., Barmet, C., Häberlin, M., Wilm, B.J., Dietrich, B.E., Schmid, T., Gross, S., Brunner, D.O., Stephan, K.E., Pruessmann, K.P., 2018. Rapid anatomical brain imaging using spiral acquisition and an expanded signal model. Neuroimage 168, 88–100.

Kasper, L., Engel, M., Heinzle, J., Mueller-Schrader, M., Jonas Reber, T.S., Barmet, C., Wilm, B.J., Stephan, K.E., Pruessmann, K.P., 2019. Advances in Spiral fMRI: A High-resolution Study with Single-shot Acquisition. BiorXiv. https://doi.org/10.1101/842179

Kasper, L., Häberlin, M., Dietrich, B.E., Gross, S., Barmet, C., Wilm, B.J., Vannesjo, S.J., Brunner, D.O., Ruff, C.C., Stephan, K.E., Pruessmann, K.P., 2014. Matched-filter acquisition for BOLD fMRI. Neuroimage 100, 145–160.

Kok, P., Bains, L.J., Van Mourik, T., Norris, D.G., De Lange, F.P., 2016. Selective activation of the deep layers of the human primary visual cortex by top-down feedback. Curr. Biol. https://doi.org/10.1016/j.cub.2015.12.038

Krämer, M., Jochimsen, T.H., Reichenbach, J.R., 2012. Functional magnetic resonance imaging using PROPELLER-EPI. Magn. Reson. Med. 68, 140–151.

Lee, G.R., Griswold, M.A., Tkach, J.A., 2010. Rapid 3D radial multi-echo functional magnetic resonance imaging. Neuroimage 52, 1428–1443.

Lustig, M., Kim, S.-J., Pauly, J.M., 2008. A fast method for designing time-optimal gradient waveforms for arbitrary k-space trajectories. IEEE Trans. Med. Imaging 27, 866–873.

Man, L.C., Pauly, J.M., Macovski, A., 1997. Multifrequency interpolation for fast off-resonance correction. Magn. Reson. Med. https://doi.org/10.1002/mrm.1910370523

Noll, D.C., Cohen, J.D., Meyer, C.H., Schneider, W., 1995. Spiral K-space MR imaging of cortical activation. J. Magn. Reson. Imaging 5, 49–56.

Nussbaum, J., Wilm, B.J., Dietrich, B.E., Pruessmann, K.P., 2018. Improved thermal modelling and prediction of gradient response using sensor placement guided by infrared photography, in: Intl. Soc. Mag. Reson. Med. Paris, p. 4210.

Pruessmann, K.P., Weiger, M., Börnert, P., Boesiger, P., 2001. Advances in sensitivity encoding with arbitrary k-space trajectories. Magn. Reson. Med. 46, 638–651.

Rahmer, J., Mazurkewitz, P., Börnert, P., Nielsen, T., 2019. Rapid acquisition of the 3D MRI gradient impulse response function using a simple phantom measurement. Magn. Reson. Med. https://doi.org/10.1002/mrm.27902

Robison, R.K., Devaraj, A., Pipe, J.G., 2010. Fast, simple gradient delay estimation for spiral MRI. Magn. Reson. Med. 63, 1683–1690.

Schleicher, A., Palomero-Gallagher, N., Morosan, P., Eickhoff, S.B., Kowalski, T., de Vos, K., Amunts, K., Zilles, K., 2005. Quantitative architectural analysis: a new approach to cortical mapping. Anat. Embryol. (Berl). 210, 373–386.

Schmitt, F., Stehling, M.K., Turner, R., 1998. Echo-Planar Imaging, Theory, Technique and Application. Springer.

Smith, S.M., Jenkinson, M., Woolrich, M.W., Beckmann, C.F., Behrens, T.E.J., Johansen-Berg, H., Bannister, P.R., De Luca, M., Drobnjak, I., Flitney, D.E., Niazy, R.K., Saunders, J., Vickers, J., Zhang, Y., De Stefano, N., Brady, J.M., Matthews, P.M., 2004. Advances in functional and structural MR image analysis and implementation as FSL. Neuroimage 23 Suppl 1, S208–19.

Spirig, Y., Graedel, N.N., Kasper, L., Miller, K.L., Frost, R., Clare, S., Pruessmann, K.P., Vannesjo, S.J., 2017. Interaction between trajectory deviations and B0 field inhomogeneity in readout-segmented EPI and spiral imaging., in: Intl. Soc. Mag. Reson. Med. p. 3917.

Stich, M., Pfaff, C., Wech, T., Slawig, A., Ruyters, G., Dewdney, A., Ringler, R., Köstler, H., 2019. Temperature-dependent gradient system response. Magn. Reson. Med. 44, 532.

Sutton, B.P., Noll, D.C., Fessler, J.A., 2003. Fast, Iterative Image Reconstruction for MRI in the Presence of Field Inhomogeneities. IEEE Trans. Med. Imaging 22, 178–188.

Vannesjo, S.J., Graedel, N.N., Kasper, L., Gross, S., Busch, J., Haeberlin, M., Barmet, C., Pruessmann, K.P., 2016. Image reconstruction using a gradient impulse response model for trajectory prediction. Magn. Reson. Med. https://doi.org/10.1002/mrm.25841

Vannesjo, S.J., Haeberlin, M., Kasper, L., Pavan, M., Wilm, B.J., Barmet, C., Pruessmann, K.P., 2013. Gradient system characterization by impulse response measurements with a dynamic field camera. Magn. Reson. Med. 69, 583–593.

Warnking, J., Dojat, M., Guérin-Dugué, A., Delon-Martin, C., Olympieff, S., Richard, N., Chéhikian, A., Segebarth, C., 2002. fMRI retinotopic mapping--step by step. Neuroimage 17, 1665–1683.

Wilm, B.J., Barmet, C., Gross, S., Kasper, L., Vannesjo, S.J., Haeberlin, M., Dietrich, B.E., Brunner, D.O., Schmid, T., Pruessmann, K.P., 2016. Single-shot spiral imaging enabled by an expanded encoding model: Demonstration in diffusion MRI. Magn. Reson. Med. 77, 83–91.

Wilm, B.J., Dietrich, B.E., Reber, J., Vannesjo, S.J., Pruessmann, K.P., 2019. Gradient response harvesting for continuous system characterization during MR sequences. IEEE Trans. Med. Imaging 1.

Woolrich, M.W., Ripley, B.D., Brady, M., Smith, S.M., 2001. Temporal autocorrelation in univariate linear modeling of FMRI data. Neuroimage. https://doi.org/10.1006/nimg.2001.0931

Wright, K.L., Hamilton, J.I., Griswold, M.A., Gulani, V., Seiberlich, N., 2014. Non-Cartesian parallel imaging reconstruction. J. Magn. Reson. Imaging 40, 1022–1040.

Yacoub, E., Harel, N., Ugurbil, K., 2008. High-field fMRI unveils orientation columns in humans. Proc. Natl. Acad. Sci. U. S. A. 105, 10607–10612.

Yang, Y., Glover, G.H., van Gelderen, P., Patel, A.C., Mattay, V.S., Frank, J.A., Duyn, J.H., 1998. A comparison of fast MR scan techniques for cerebral activation studies at 1.5 tesla. Magn. Reson. Med. 39, 61–67.

Zhang, Y., 0001, M.B., Smith, S.M., 2001. Segmentation of Brain MR Images through a Hidden Markov Random Field Model and the Expectation Maximization Algorithm. IEEE Trans. Med. Imaging 20, 45–57.

Zilles, K., Amunts, K., 2010. Centenary of Brodmann’s map--conception and fate. Nat. Rev. Neurosci. 11, 139–145.

